# CWHM-974 is a fluphenazine derivative with improved antifungal activity against *Candida albicans* due to reduced susceptibility to multidrug transporter-mediated resistance mechanisms

**DOI:** 10.1101/2023.05.01.538946

**Authors:** Aracely Miron-Ocampo, Sarah R. Beattie, Soumitra Guin, Thomas Conway, Marvin J. Meyers, W. Scott Moye-Rowley, Damian J. Krysan

## Abstract

Multidrug resistance (MDR) transporters such as ATP Binding Cassette (ABC) and Major Facilitator Superfamily (MFS) proteins are important mediators of antifungal drug resistance, particularly with respect to azole class drugs. Consequently, identifying molecules that are not susceptible to this mechanism of resistance is an important goal for new antifungal drug discovery. As part of a project to optimize the antifungal activity of clinically used phenothiazines, we synthesized a fluphenazine derivative (CWHM-974) with 8-fold higher activity against *Candida* spp. compared to the fluphenazine and with activity against *Candida* spp. with reduced fluconazole susceptibility due to increased multidrug resistance transporters. Here, we show that the improved *C. albicans* activity is because fluphenazine induces its own resistance by triggering expression of CDR transporters while CWHM-974 induces expression but does not appear to be a substrate for the transporters or is insensitive to their effects through other mechanisms. We also found that fluphenazine and CWHM-974 are antagonistic with fluconazole in *C. albicans* but not in *C. glabrata*, despite inducing *CDR1* expression to high levels. Overall, CWHM-974 represents a unique example of a medicinal chemistry-based conversion of chemical scaffold from MDR-sensitive to MDR-resistant and, hence, active against fungi that have developed resistance to clinically used antifungals such as the azoles.

## Introduction

Phenothiazines are one of the oldest classes of drugs with derivatives of this class being used to treat a wide range of diseases from parasitic infections to nausea to psychosis (1). One of the features of the phenothiazine class of molecules is that it readily crosses the blood-brain-barrier (1). Accordingly, most of the phenothiazine-derived drugs currently in use are for the treatment of CNS-related conditions. Clinically used phenothiazines have also been studied for repurposing to other indications with anti-cancer and anti-infective applications being amongst the most widely investigated (2).

The potential of phenothiazines such as trifluoperazine, thioridazine and fluphenazine as antifungal agents has been investigated for many years (3), including work from our group (4). An important dose-limiting toxicity of these drugs is their antipsychotic and sedative CNS activity. As part of our efforts in this area, we reported a fluphenazine derivative, CWHM-974 (Fig. 1), with increased antifungal activity against fungal pathogens such as *C. albicans* and *C. neoformans* as well as reduced affinity for dopamine and histamine receptors (4).

**Figure 1.**
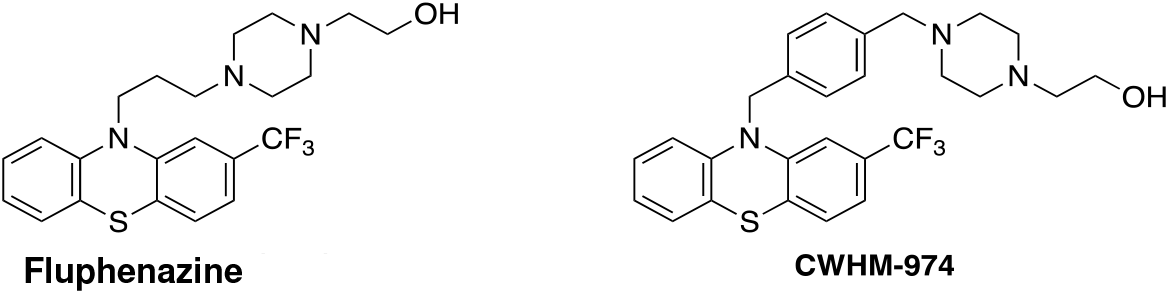
Chemical structures of fluphenazine and CWHM-974.

Fluphenazine, the parent drug of CWHM-974, has poor activity against *C. albicans* with a minimum inhibitory concentration (MIC) of 64 μg/mL while CWHM-974 is 8-fold more active (MIC; 8 μg/mL). Fluphenazine is a strong inducer of ATP-Binding Cassette (ABC) and Major Facilitator Superfamily (MFS) proteins such as *CDR1* and *MDR1* in *C. albicans* and, as such, is frequently used to study *C. albicans* the regulation of multidrug transporter expression (5, 6). One of the important features of CWHM-974 is that it is active against a *C. albicans* strain that is fluconazole-resistant due to increased expression of *CDR1* and *MDR1* (4). We, therefore, were interested to determine if the increased anti-*C. albicans* activity of CWHM-974 relative to fluphenazine was because of altered interactions with the multidrug transporters or their regulators. Specific questions were: 1) does CWHM-974 induce pump expression? 2) Is CWHM-974 less susceptible to efflux pump-mediated resistance? 3) does CWHM-974 inhibit efflux pump function?

Here, we show that both CWHM-974 and fluphenazine induce the ABC transporters *CDR1* and *CDR2* and that CWHM-974 is a more potent inducer than fluphenazine. We also found that fluphenazine is susceptible to multi-drug transporter mediated resistance while CWHM-974 is not. Both fluphenazine and CWHM-974 antagonize fluconazole in a Tac1/Cdr-dependent manner, indicating that CWHM-974 is unlikely to inhibit transporter function. To our knowledge, the CWHM-974 is one of the few examples of a molecule in which relatively small structural modifications significantly reduced susceptibility to multidrug transporter-mediated resistance.

## Results

### Fluphenazine and CWHM-974 induce distinct patterns of efflux drug expression

Fluphenazine is a well-studied inducer of *CDR1* expression in *C. albicans* (6, 7). We confirmed those observations (Fig. 2A) and found that CWHM-974 also induces robust *CDR1* expression. Concentrations of the two molecules for these experiments were normalized to their respective MIC value and, thus, CWHM-974 leads to ∼2.5-fold higher *CDR1* expression compared to fluphenazine. Both molecules also induce the expression of *CDR2* (Fig. 2A) but only CWHM-974 induces *MDR1* expression. Thus, CWHM-974 appears to be a more potent and more general inducer of drug ABC/MDR expression than the fluphenazine parent compound.

**Figure 2.**
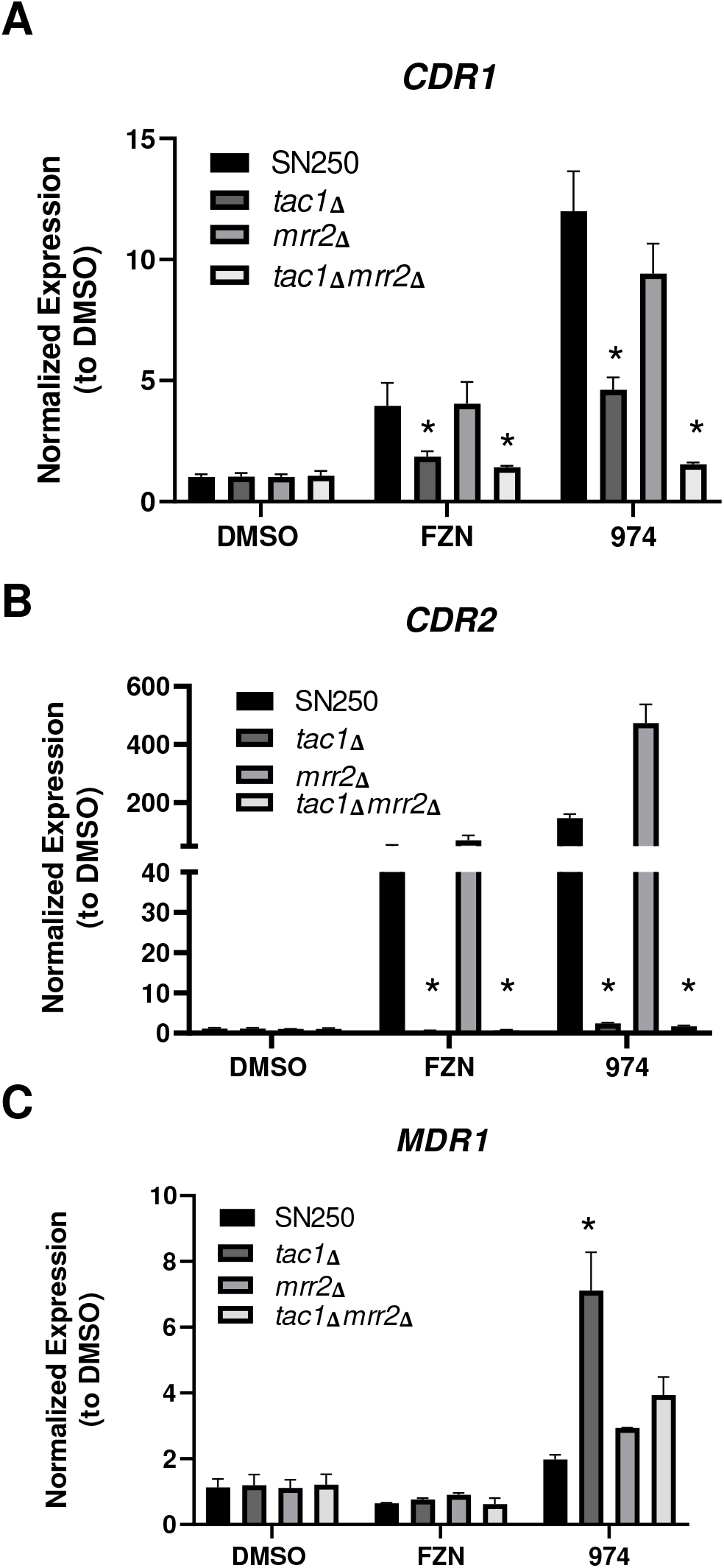
Fluphenazine and CWHM-974 induce distinct *CDR1/2* and *MDR1* transporter expression patterns. The expression of the indicated genes was assayed using quantitative RT-PCR of RNA isolated from the indicated strains after 2 hr exposure of logarithmic phase cells to fluphenazine (FZN, 50 μg/mL), CWHM-974 (6.25 μg/mL), DMSO carrier alone (1%). The expression levels were normalized to DMSO treated cells and data are mean fold-change from triplicate experiments with the error bars indicating the standard deviation. Statistical significance was determined using 2-way ANOVA with Dunnett’s correction for multiple comparisons and defined as an adjusted p value <0.05. Statistically significant differences from the SN250 reference are indicated by an asterisk (*). Effect of Tac1 and Mrr2 on expression of *CDR1* (**A**), *CDR2* (**B**), and *MDR1* (**C**).

Previous work has shown that fluphenazine-induced expression of *CDR1* and *CDR2* is dependent on the transcription factor Tac1 (6, 7). As shown in Fig. 2A, deletion of *TAC1* reduces both *CDR1* and *CDR2* expression in cells exposed to either fluphenazine or CWHM-974. However, CWHM-974-induced *CDR1* expression remains significantly elevated relative to untreated cells in the *tac1*ΔΔ strain. Gain-of-function alleles of *MRR1* also increase expression of *CDR1* (8) and, therefore, we asked if Mrr2 contributes to CWHM-974 induced expression. To test this, we constructed a *tac1*ΔΔ *mrr2*ΔΔ double mutant and compared CWHM-974-induced expression in this strain to the two single mutants. In the presence of fluphenazine (Fig. 2A), *CDR1* expression did not differ from WT in the *mrr2*ΔΔ mutant and the *tac1*ΔΔ *mrr2*ΔΔ mutant did not differ from *tac1*ΔΔ. In contrast, *CDR1* expression was modestly reduced in the *mrr2*ΔΔ mutant exposed to CWHM-974 while it was returned to the level of untreated cells in the *tac1*ΔΔ *mrr2*ΔΔ double mutant. These genetic interaction data strongly indicate that CWHM-974 induces *CDR1* expression through both Tac1 and Mrr2 while fluphenazine activates expression almost exclusively through Tac1.

Tac1 is also required for *CDR2* induction for both fluphenazine and CWHM-974 (6, 7). Curiously, while the *mrr2*Δ mutant has no effect on fluphenazine-induced *CDR2* expression (Fig. 2B), it dramatically accentuates CWHM-974 induced *CDR2* expression. The increased *CDR2* expression in the *mrr2*ΔΔ mutant is completely abolished by the deletion of *TAC1*. This observation suggests that Mrr2 negatively modulates CWHM-974 activation of Tac1 either directly or indirectly.

A similar phenomenon was observed with CWHM-974 induction of *MDR1* expression (Fig. 2C). In this case, however, deletion of *TAC1* leads to a 4-fold increase in *MDR1* expression relative to the wildtype strain. Deletion of *MRR2* also increases CWHM-974-induced expression of *MDR1*. The level of *MDR1* expression induced in the *tac1*ΔΔ *mrr2*ΔΔ double mutant expression is, however, slightly reduced relative the *tac1*ΔΔ, suggesting that these two transcription factors may make some contribution to the positive regulation of *MDR1*. However, the expression is still 2-fold higher than wildtype indicating that other factors must regulate *MDR1* expression. Taken together, these data indicate that the fluphenazine derivatives induce multiple efflux pumps through a complex inter-related network of regulators. Our CWHM-974 data also indicate compensatory interplay between *CDR1/2* expression and *MDR1* expression that is mediated by Mrr1/Tac1 and an unidentified regulator of *MDR1* expression. Finally, these results indicate that the improvement in anti-*C. albicans* activity of CWHM-974 is not due to reduced induction of ABC/MFS multidrug transporter proteins.

### The antifungal activity of CWHM-974 is required for induction of *CDR1* expression

The antifungal activity of fluphenazines and other phenothiazines is due in part to inhibition of calmodulin-dependent processes (4, 8). In contrast, the molecular mechanism by which these molecules induce multidrug resistance transporter expression is not clear. Fluphenazine has reduced antifungal activity and reduced potency with respect to *CDR1* expression relative to CWHM-974, suggesting that there may be a correlation between these two properties. We, therefore, asked if the antifungal activity of the fluphenazine derivatives was required to induce *CDR1* expression. Oxidation of the S atom in the phenothiazine core structure blocks the ability of the molecule to bind calmodulin (8) and we previously showed that oxidation of the phenothiazine antipsychotic thioridazine abolished its antifungal activity (4). We synthesized the sulfoxide of CWHM-974 (894) and found that it had no antifungal activity at concentrations up to the limit of its solubility (Fig. 3A). 894 also did not induce *CDR1* expression at the highest soluble concentrations (Fig.3B). This observation suggests that antifungal activity is required for fluphenazine-induced *CDR1* expression and further supports the notion that the potency of antifungal activity correlates with the extent of *CDR1* expression in the fluphenazine series. One explanation for these observations is that *CDR1* expression is in response to the physiologic changes caused by inhibition of the antifungal target of fluphenazine/CWHM-974. A second possibility is that the receptor that recognizes multidrug transporter inducing molecules shares structural features with the antifungal target of the fluphenazines and, therefore, is subject to a similar structure-activity relationship.

**Figure 3.**
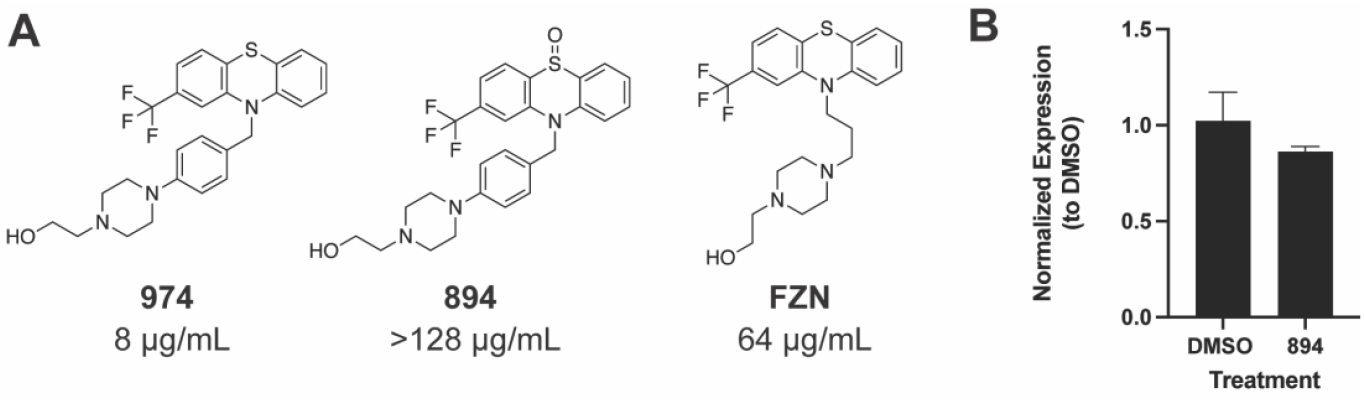
*S*-oxidation of CWHM-974 eliminates antifungal activity and ability to induce *CDR1*. **A**. Chemical structures of CWHM-974, CWHM-894, and fluphenazine (FZN) along with their MIC values against *C. albicans*. **B**. *CDR1* expression of SN250 cells exposed to CWHM-974 (128 μg/mL) compared to DMSO carrier solvent alone (1%).

### The antifungal activity of fluphenazine but not CWHM-974 is modulated by the interactions of Tac1, Mrr2, and Cdr1

Next, we asked whether the mutants of the Tac1/Mrr2/Cdr system contributed to the antifungal susceptibility of *C. albicans* to fluphenazine or CWHM-974. As shown in Table 1, single deletion mutants of *TAC1, MRR2*, and *CDR1* did not show significant difference in susceptibility to either fluphenazine or CWHM-974 under modified CLSI microbroth dilution conditions (temperature 37°C rather than 35°C). Double and triple mutants involving Tac1, Mrr2, and Cdr1 had no significant effect on the MIC of CWHM-974. In contrast, the *tac1*ΔΔ *cdr1*ΔΔ mutant was 4-fold more susceptible to fluphenazine. The fluphenazine MIC of neither the *tac1*ΔΔ *mrr2*ΔΔ nor the *mrr2*ΔΔ *cdr1*ΔΔ mutant was significantly different than WT but the *tac1*ΔΔ *mrr2*ΔΔ *cdr1*ΔΔ triple mutant had the lowest fluphenazine MIC (4 μg/mL; 16-fold reduction from WT). These data indicate that fluphenazine is highly sensitive to the Tac1-Mrr2-Cdr multidrug transporter while CWHM-974 is not. This strongly suggests that the structural modifications made to CWHM-974 block the ability of these transporters to modify their activity, particularly since the modification increase its potency as an inducer of this drug transporter system.

**Table 1.**
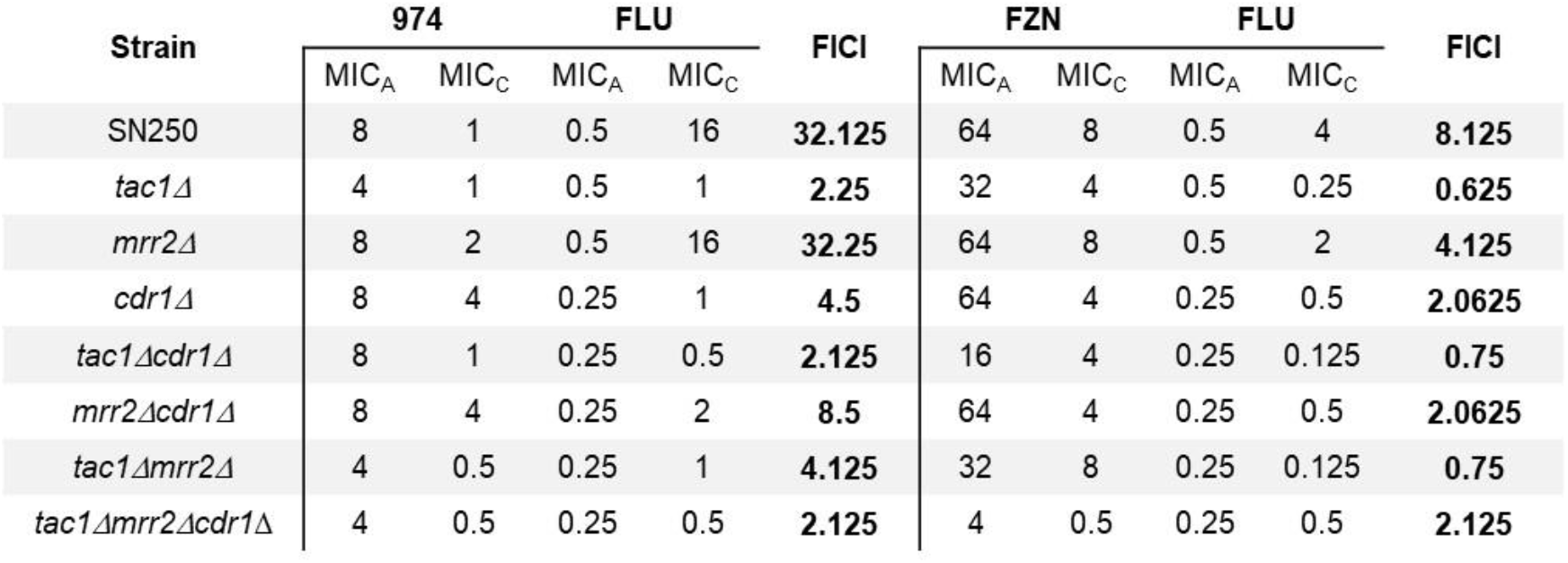

| Strain | 974 |  | FLU |  | FICI | FZN |  | FLU |  | FICI |
| --- | --- | --- | --- | --- | --- | --- | --- | --- | --- | --- |
|  | MIC<sub>A</sub> | MIC<sub>C</sub> | MIC<sub>A</sub> | MIC<sub>C</sub> |  | MIC<sub>A</sub> | MIC<sub>C</sub> | MIC<sub>A</sub> | MIC<sub>C</sub> |  |
| SN250 | 8 | 1 | 0.5 | 16 | 32.125 | 64 | 8 | 0.5 | 4 | 8.125 |
| *tac1Δ* | 4 | 1 | 0.5 | 1 | 2.25 | 32 | 4 | 0.5 | 0.25 | 0.625 |
| *mrr2Δ* | 8 | 2 | 0.5 | 16 | 32.25 | 64 | 8 | 0.5 | 2 | 4.125 |
| *cdr1Δ* | 8 | 4 | 0.25 | 1 | 4.5 | 64 | 4 | 0.25 | 0.5 | 2.0625 |
| *tac1Δcdr1Δ* | 8 | 1 | 0.25 | 0.5 | 2.125 | 16 | 4 | 0.25 | 0.125 | 0.75 |
| *mrr2Δcdr1Δ* | 8 | 4 | 0.25 | 2 | 8.5 | 64 | 4 | 0.25 | 0.5 | 2.0625 |
| *tac1Δmrr2Δ* | 4 | 0.5 | 0.25 | 1 | 4.125 | 32 | 8 | 0.25 | 0.125 | 0.75 |
| *tac1Δmrr2Δcdr1Δ* | 4 | 0.5 | 0.25 | 0.5 | 2.125 | 4 | 0.5 | 0.25 | 0.5 | 2.125 |

### Fluphenazine and CWHM-974 induce Tac1-Cdr1-dependent fluconazole antagonism in *C. albicans*

The combination of fluphenazine and fluconazole is antagonistic against *C. albicans* (9, 10), presumably by induction of *CDR1/2*, although that has not been experimentally confirmed. To test this hypothesis further, we first asked whether the combination of fluconazole and fluphenazine induced *CDR1* in an additive or synergistic manner. As shown in Fig. 4A, expression of *CDR1* is higher in *C. albicans* cells treated with fluconazole and either fluphenazine or CWHM-974 than in cells treated with either drug alone. The expression of *CDR1* in the presence of both fluconazole and fluphenazine is equal to the product of the expression levels of each drug alone, indicating an additive effect. The expression of *CDR1* is, however, less than the product of the expression observed in cells treated with either fluconazole or alone CWHM-974, indicating that the expression may be reaching saturation.

**Figure 4.**
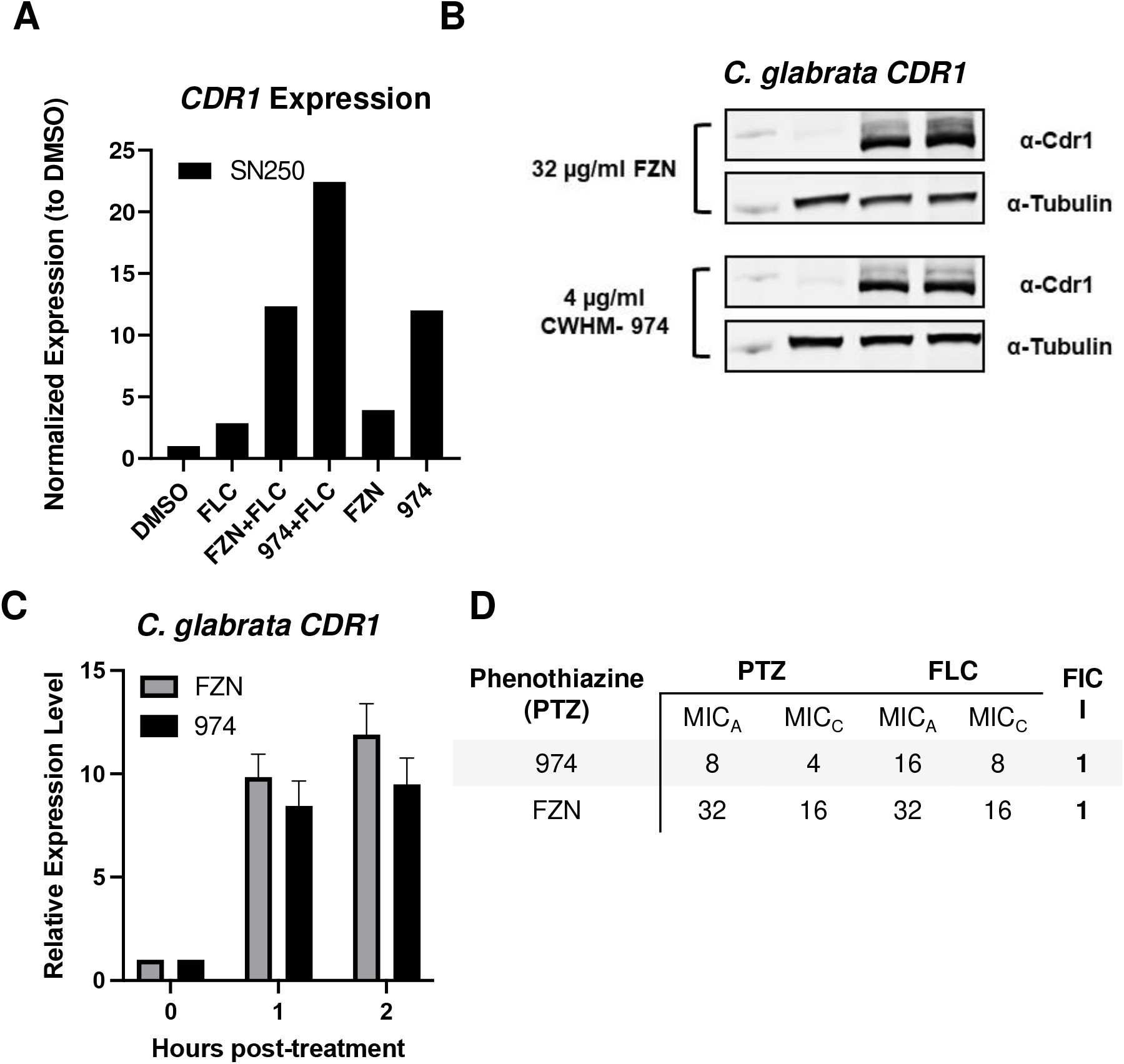
Fluphenazine and CWHM-974 induce *CDR1* expression but are not antagonistic. **A**. Fluconazole and fluphenazines combine to increase *CDR1* expression relative to single drug treatments. SN250 cells were treated with both fluconazole and either FZN or CWHM-974 at concentrations 2-fold below their respective FIC values (Table 1) for 2 hr before being processed for quantitative RT-PCR. All treatments showed statistically significant changes from untreated SN250 cells. **B**. Representative western blot of Cdr1 expression in the presence of the indicated concentrations of FZN and CWHM-974. **C**. Quantitative analysis with bars indicating the mean and error bars the standard deviation of duplicate western blots. The treated cells are statistically different than untreated but there is no statistical difference between FZN and CWHM-974. **D**. FIC analysis of the interaction of FZN and CWHM-974 with fluconazole in *C. glabrata*.

This observation is consistent with a model in which all three molecules induce expression of *CDR1* through a common mechanism or pathway.

Consistent with previous work (9, 10), both fluphenazine and CWHM-974 are antagonistic with fluconazole in *C. albicans* by checkerboard analysis (Table 1). Interestingly, both fluphenazine and CWHM-974 strongly inducing the expression of *CDR1* (Fig. 4B&C). However, there is no antagonism between either fluphenazine or CWHM-974 and fluconazole in *C. glabrata* (Fig. 4D). In *C. albicans* (Table 1), CWHM-974 treatment causes a 32-fold increase in the fluconazole MIC while the MIC of fluconazole is 8-fold higher in combination with fluphenazine, suggesting that the higher expression of *CDR1* in the presence of CWHM-974 and fluconazole translates to lower susceptibility. In the presence of fluconazole, the MIC of the fluphenazine and CWHM-974 is reduced 8-fold, suggesting that inhibition of Erg11 increases the activity of the phenothiazines. Thus, the antagonism, as analyzed by this checkerboard approach, is driven solely by the increase in fluconazole MIC.

Next, we determined the effect of *tac1*ΔΔ, *cdr1*ΔΔ, and *tac1*ΔΔ *cdr1*ΔΔ mutations on the increase in fluconazole MIC induced by the fluphenazine derivatives. As shown in Table 1, deletion of either *TAC1* or *CDR1* return the combination fluconazole MIC to within a 2-fold dilution of that observed for fluconazole alone. The strain lacking both *TAC1* and *CDR1* has a combination fluconazole MIC identical to that of wild type treated with fluconazole alone. Thus, the induction of antagonism by fluphenazine and CWHM-974 can be explained by activation of the Tac1-Cdr1 axis. Phenothiazines have been shown to inhibit efflux pump function in other settings (10), raising the possibility that the increased anti-*C. albicans* activity of CWHM-974 may be due to its ability to inhibit Cdr1. If that were the case, then CWHM-974 would induce CDR1 but there would be no antagonism with fluconazole. Consequently, these data are inconsistent with multidrug transport protein inhibition contributing to the improved activity of CWHM-974 against *C. albicans* relative to fluphenazine.

## Discussion

Multidrug transport mediated resistance is one of the most clinically important mechanisms of antifungal drug resistance because it reduces susceptibility to the azole class of molecules in multiple fungal pathogens including *Candida* spp and *Cryptococcus* spp (12). Identifying new drugs that are not susceptible to these mechanisms is critical to improving the overall efficacy of antifungal chemotherapy. A wide range of non-antifungal drugs and bioactive small molecules induce the expression of ABC and MFS transporters in fungi (9, 13). Of these one of the most widely studied is fluphenazine, particularly with respect to ABC transporter expression in *C. albicans*. As part of a repurposing project aimed at the medicinal chemistry-based optimization of phenothiazine antifungal activity (4), we synthesized a derivative of fluphenazine (CWHM-974) with reduced affinity for dopamine and histamine receptors. CWHM-974 also showed improved activity against *C. albicans* relative to the parent fluphenazine. This improved activity was also observed for strains that are fluconazole resistant due to increased expression of CDR and MFS transporters, suggesting that CWHM-974 may not be susceptible to ABC/MFS mediated resistance mechanisms.

The genetic analysis described provides strong support for this hypothesis. First, we have confirmed that fluphenazine induces the expression of *CDR1* and *CDR2* and that the increased expression of these transporters is responsible for its poor antifungal activity against *C. albicans*. Second, CWHM-974 is a more potent inducer of *CDR1/2* but mutants lacking these genes or with dramatically reduced expression of *CDR1/2* show no change in susceptibility compared to WT. Third, CWHM-974 and fluphenazine potently induce Tac1-Cdr1-mediated fluconazole antagonism, indicating that CWHM-974 is unlikely to interfere indirectly or directly with the function of this resistance pathway to a significant extent. We are unaware of another example whereby structural modification of an antifungal small molecule that is susceptible to CDR/MFS mediated resistance leads to abrogation of that resistance.

At this point, it is not clear how replacement of the alkyl chain linking the phenothiazine core to the piperazine moiety of fluphenazine with a larger, benzyl linker causes this dramatic change in sensitivity to the CDR resistance mechanism. The simplest explanation is that, as proposed in the literature (9, 13), fluphenazine is a substrate for the efflux system while the benzyl-modified CWHM-974 is not. Additional work will be required to identify the structural and biochemical basis for the ability of CWHM-974 to evade CDR-mediated resistance mechanisms.

Our genetic interaction analysis of the Tac1-Mrr2-CDR system with respect to fluconazole susceptibility has led to observations that warrant emphasis. First, deletion of multiple regulators, with or without deletion of *CDR1*, had almost no effect on the susceptibility of those strains to fluconazole. Second, and consistent with this finding, fluconazole alone is a relatively weak inducer of *CDR1* expression. Consequently, fluconazole is not a sufficiently strong inducer of *CDR1* expression to mediate its own resistance. Fluphenazine, on the other hand, induces sufficient *CDR1* and *CDR2* expression to mediate its own resistance. In addition, fluphenazine resistance can be mediated by either *CDR1* or *CDR2* or possibly other transporters since the *tac1*ΔΔ *mrr1*ΔΔ *cdr1*ΔΔ mutant is more susceptible than the single mutants or the *tac1*ΔΔ *mrr1*ΔΔ double mutant. In contrast to fluphenazine-induced, fluphenazine resistance, fluphenazine/CWHM-974-induced, fluconazole resistance is completely derived from Tac1-dependent expression of *CDR1*.

Finally, it is worth noting that neither fluphenazine nor CWHM-974 show antagonism with fluconazole in *C. glabrata* despite inducing a 10-fold increase in *CDR1* expression (Fig. 4B-D). *C. glabrata* strains with elevated *CDR1* expression by virtue of Pdr1 gain-of-function mutants are resistant to fluconazole compared to strains with WT alleles (14). Indeed, *pdr1*^*R376W*^ has a nearly identical 10-fold increase Cdr1 protein levels as that observed with fluphenazine and CWHM-974 exposure (14). Consistent with that increased expression, the *pdr1*^*R376W*^ mutant has reduced susceptibility to fluconazole compared to the WT strain (14). Based on the similar levels of *CDR1* expression, one would expect fluphenazine and CWHM-974 to induce antagonism. The lack of antagonism between the fluphenazines and fluconazole, however, suggests that fluphenazine and CWHM-974 have additional effects on fluconazole susceptibility that counter the increased expression of *CDR1*.

In summary, we have found that the fluphenazine derivative CWHM-974 displays increased activity against *C. albicans* because it is less susceptible to CDR/MFS mechanisms of resistance. Efforts to further optimize the antifungal activity and therapeutic index of this promising, repurposed scaffold are ongoing.

## Materials and methods

### Strains, media and reagents

All yeast strains were in the SN background and maintained on yeast peptone (2%) dextrose (YPD) plates after recovery from 25% glycerol stocks stored at - 80°C. Transcription factor mutants *tac1*ΔΔ and *mrr2*ΔΔ were obtained from the Homann deletion collection provided by the Fungal Genetics Stock Center (15). The *cdr1*ΔΔ, *tac1*ΔΔ *mrr2*Δ and *tac1*ΔΔ *mrr2*ΔΔ *cdr1*ΔΔ mutants (see Table S1 for strains) were generated using transient CRISPR methods described by Huang et al. The mutants were confirmed using PCR analysis of the marker junctions and lacked the targeted ORF by PCR analysis; see Table S2 for primers used for these analyses. Yeast media were prepared using recipes described in Homann et al (15). Fluconazole and fluphenazine were obtained from Sigma-Aldrich. CWHM-974 was synthesized as previously reported and was greater than 95% pure.

### Quantitative RT-PCR analysis

The indicated strains were pre-cultured overnight in YPD at 30°C and then back-diluted to a density between 0.9 and 0.15 OD_600_. The resulting cultures were incubated for 3 hr at 30°C (OD_600_ ∼ 0.3-0.4). The strains were exposed to DMSO carrier solvent (final concentration 1%), fluphenazine (50 μg/mL), CWHM-974 (6.25 μg/ml) or fluconazole (16 μg/mL) for 2 hr at 30oC.

The cells were harvested and frozen at -80°C prior to isolation of RNA using the MasterPure Yeast RNA purification kit. 500ng of RNA was used for cDNA synthesis with iScript cDNA synthesis kit. The resulting cDNA was diluted 1:5 processed using the qRT-PCR with iQ SYBR Green Supermix with 0.20 μM primers (See Table S2 for PCR primers) qRT-PCR was performed on the BioRad CFX Connect using a 3-step amplification with 54ºC annealing temperature and melt curve analysis. Gene expression was normalized to *ACT1* and analyzed using the ΔΔ Ct method. The experiments were performed in triplicate and differences in expression as analyzed by 2-way ANOVA with Dunnett’s correction for multiple comparisons with significance limit defined as an adjusted p value < 0.05.

### Minimum inhibitory concentration and fractional inhibitory concentration determinations

Minimum inhibitory concentrations (MIC) and fractional inhibitory concentrations (FIC) were determined using modification of the CLSI conditions. All yeasts were cultured overnight in 3 mL YPD at 30°C, then washed twice in sterile PBS. Two-fold serial dilutions of each compound were prepared in RMPI+MOPS pH 7 (Gibco RPMI 1640 with L-glutamine and 0.165M MOPS) with DMSO as carrier solvent (final concentration 1%), then 1 × 10^3^ cells were added per well. Plates were incubated at 37°C for 24 h and the MIC/FIC values were determined visually as the lowest concentration with a clear well (fluphenazine and CWHM-974) or with a distinct reduction from DMSO only well. Assays were performed in duplicate or triplicate.

### Synthesis of 10-[(p-{[4-(2-hydroxyethyl)-1-piperazinyl]methyl}phenyl)methyl]-2-(trifluoromethyl)-5λ^4^-5-phenothiazinone (SLU-10894)

To a stirred solution of 2-(4-(4-((2-(trifluoromethyl)-10*H*-phenothiazin-10-yl)methyl)benzyl)piperazin-1-yl)ethanol (CWHM-974; 0.1 mmol) in MeOH (1 mL) was added a solution of NaIO_4_ (0.1 mmol) in MeOH containing one drop of conc. HCl was added dropwise. The reaction mixture was then stirred at room temperature for 15-20 min. The reaction turned dark red. It was then quenched by the addition of a few drops of water and extracted with EtOAc (2×25 mL). The combined organic layer was then dried over anhydrous Na_2_SO_4_ and concentrated under reduced pressure to afford the crude. The crude was then purified by reverse-phase HPLC (5-100 % CH_3_CN/H_2_O + 0.1 % formic acid). The pure fractions were then lyophilized to afford the titled compound as a white solid (11.2 mg, 22%). ^1^H NMR (400 MHz, DMSO) δ 8.27 (d, *J* = 7.7 Hz, 1H), 8.07 (d, *J* = 6.2 Hz, 1H), 7.67 (d, *J* = 8.7 Hz, 1H), 7.65 – 7.55 (m, 2H), 7.49 (d, *J* = 8.4 Hz, 1H), 7.40-7.34 (m, 1H), 7.24 (d, *J* = 7.9 Hz, 2H), 7.11 (d, *J* = 7.9 Hz, 2H), 5.76 (s, 2H), 4.33 (br s, 1H), 3.46 (s, 2H), 3.40 (s, 2H), 2.44 – 2.21 (m, 10H); HPLC purity >96%. LCMS m/z 516 [M+H]^+^. HRMS (ESI) m/z: [M + H]^+^ Calcd for C_27_H_28_F_3_N_3_O_2_S 516.1932, found 516.1933.

### Western blot analysis of *C. glabrata* Cdr1

*C. glabrata* ATCC 2001 was used for the analysis of the induction of Cdr1 expression under different drug conditions (14). Cultures of *C. glabrata* were grown in YPD (1% yeast extract, 2% peptone, 2% glucose) medium at 30°C. Overnight cultures were diluted to an OD_600_ of 0.2 in fresh YPD, grown at 30°C to an OD_600_ of 0.4, split between three flasks, and treated separately with either fluconazole (16 μg/ml), fluphenazine (32 μg/ml), or 974 (4 μg/ml). Cell samples were acquired immediately prior to splitting the initial cultures and at the 1 and 2 hours after drug treatment. Proteins were extracted as previously described (Shahi et al., 2010), resuspended in urea sample loading buffer (8M urea, 1% 2-mercaptoethanol, 40 mM Tris-HCL pH 8.0, 5% SDS, bromophenol blue), and incubated at room temperature overnight. Protein extracts were resolved in an ExpressPlus 4-12% gradient gel (GenScript), electroblotted to a nitrocellulose membrane, blocked with 5% nonfat dry milk, and probed with an anti-Cdr1 polyclonal antibody (Vu et al., 2019) and an anti-alpha-tubulin monoclonal antibody (12G10; Developmental Studies Hybridoma Bank at the University of Iowa). For detection and quantification of blotted proteins, secondary Li-Cor antibodies, IRD dye 680RD goat anti-rabbit and IRD dye 800LT goat anti-mouse, were used in combination with the Li-Cor infrared imaging system (application software version 3.0) and Image Studio Lite software (Li-Cor). Cdr1 protein levels were normalized to tubulin and compared to the pretreatment condition. Measurements represent the result of two technical replicates each from two biological replicates.

## Acknowledgements

This work was funded in part by NIH grants: R21AI164578 (DJK and MVM), F32AI145160 (SRB) and R01AI52494 (WSMR). We thank Rohan S. Wakade (Iowa) for assistance with the generation of *C. albicans* mutants.

## Supplementary Material

**Table S1. Strain Table**

**Table S2. Oligonucleotide Table**

